# Deciphering Phage-Host Specificity Based on the Association of Phage Depolymerases and Bacterial Surface Glycan with Deep Learning

**DOI:** 10.1101/2023.06.16.545366

**Authors:** Yiyan Yang, Keith Dufault-Thompson, Wei Yan, Tian Cai, Lei Xie, Xiaofang Jiang

## Abstract

Phage tailspike proteins are depolymerases that target diverse bacterial surface glycans with high specificity, determining the host-specificity of numerous phages. To address the challenge of identifying tailspike proteins due to their sequence diversity, we developed SpikeHunter, an approach based on the ESM-2 protein language model. Using SpikeHunter, we successfully identified 231,965 tailspike proteins from a dataset comprising 8,434,494 prophages found within 165,365 genomes of five common pathogens. Among these proteins, 143,035 tailspike proteins displayed strong associations with serotypes. Moreover, we observed highly similar tailspike proteins in species that share closely related serotypes. We found extensive domain swapping in all five species, with the C-terminal domain being significantly associated with host serotype highlighting its role in host range determination. Our study presents a comprehensive cross-species analysis of tailspike protein to serotype associations, providing insights applicable to phage therapy and biotechnology.

## Introduction

Understanding the host specificity of bacteriophages is a crucial part of developing phage therapies for bacterial infections and applying phage proteins in biotechnology. Extensive work has been done to understand what bacteria phages can infect and what proteins are involved in this process^1, 2^. Experimental determination of bacteriophage host range can be slow and challenging, often being limited to small collections of bacterial and phage strains^1, 2^. Computational approaches have the benefit of being able to predict host range based on genomic information^3^, but these methods are often limited in taxonomic resolution and sensitivity. A computational approach that utilizes large-scale, readily available genomic data and is based on the underlying mechanisms of phage host range determination has the potential to provide improved sensitivity and predictive power, serving as a valuable tool for guiding future phage therapy endeavors.

Tailspike proteins, which function as both depolymerases and receptor-binding proteins, play a crucial role in determining phage host range. These proteins specifically recognize types of bacterial cell surface glycans, including Capsular Polysaccharides (CPS) and Lipopolysaccharides (LPS), allowing the phage to attach to and degrade the glycans as part of the infection process^4^. In phages with tailspike proteins, they play a major role in determining the host range of the phage due to bacterial strains expressing different types of surface glycans including K-antigen CPS glycans, O-antigen LPS glycans, and OC-antigens^5–7^. Varieties of these surface glycan antigens differ in their sugar composition and the linkages connecting those sugars, providing a diverse range of sugar motifs for tailspike proteins to recognize on different hosts. Understanding the association of these tailspike proteins with specific bacterial serotypes, the types of glycans on a bacteria’s surface, can help streamline the process of selecting effective phages for phage therapy, one of the major bottlenecks in the broader use of this treatment^8^. Phage tailspike proteins also have potential applications as antimicrobials, having been shown to sensitize resistant strains to serum killing^9^ and degrade biofilms^10^. These applications and the engineering of phages to target new bacterial strains^11^ or avoid bacterial resistance^12^, would significantly benefit from a better understanding of phage-host specificity.

Given their central role in phage-host interactions, many studies have attempted to understand the associations of tailspike proteins with bacterial serotypes. Recent work has demonstrated that phage host range is typically limited and has provided valuable insights into the importance of the phage tailspike proteins in determining this range^2^. Similarly, other studies related to *Ackermannviridae* and *Escherichia* viruses have demonstrated that tailspike proteins are tightly associated with host serotype and that recombination of tailspike protein domains may be a key driver in tailspike protein evolution^2, 13, 14^. While these studies have provided valuable examples of the association of tailspike proteins with specific serotypes, their scale and scope, often focusing on single bacterial species and fewer than a hundred phages, limits their generalizability and predictive power, especially in the context of bioengineering and phage therapy.

In this study we present a large-scale multi-species analysis of tailspike protein associations with bacterial serotypes. Using a deep-learning classifier based on a protein language model, we identified prophage-encoded tailspike proteins in an extensive dataset of genomes from five common human pathogens. These tailspike proteins were combined with predicted serotypes for the bacteria to generate a large-scale dataset of tailspike protein-serotype associations that captures the broad diversity seen in these proteins. By analyzing these associations, we identified many strongly associated tailspike-serotype pairs, examples of cross-species tailspike associations, and evidence of extensive domain swapping between tailspike proteins. These results benefit from the large scale of the analysis, covering thousands of strains across multiple bacterial species. This provides an expansive dataset of tailspike proteins that can be used to make informed predictions about the host range of other phages and guide future applications of phages and phage proteins.

## Results

### A deep learning powered model for detecting sequence-divergent tailspike proteins

To investigate the association between tailspike proteins and bacterial surface glycan antigens, we developed the SpikeHunter deep-learning model to identify tailspike proteins (**Fig.1a**). SpikeHunter was built on the ESM2 large protein language model^15^ to embed a protein sequence into a representative vector and predict the probability of that protein being a tailspike protein using a fully connected three-layer neural network (**Fig.1b**). A reference set of 1,913 tailspike protein sequences and 201,994 non-tailspike protein sequences was curated from the INPHARED database^16^. The labeled protein sequences were then partitioned into training, validation, and testing datasets at a ratio of 3:1:1. This partition was conducted to preserve an equivalent proportion of positive samples across each set and also to ensure that no sequence within a particular set displayed over 30% identity with a sequence in another set.

**Fig.1.**
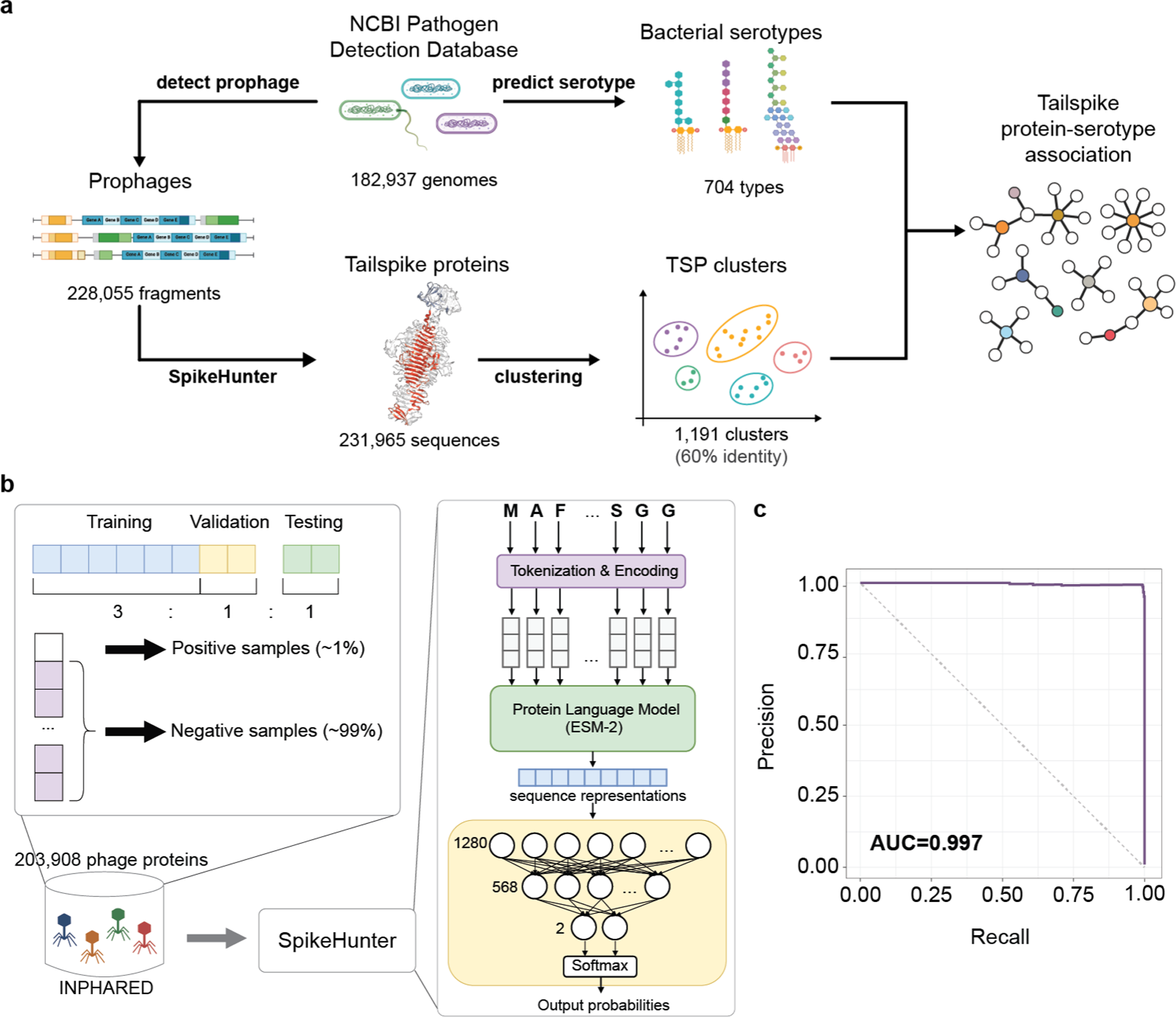
Development of the SpikeHunter deep-learning model. a, Diagram showing the project workflow for the identification of tailspike protein and serotype associations. b, Organization of the training data for the SpikeHunter model and deep-learning model architecture for SpikeHunter. C, The precision-recall curve with an area under the curve of 0.990 for SpikeHunter evaluated on a testing dataset consisting of 40,238 phage proteins.

Early stopping was adopted to stop training once the model performance was no longer improved on the validation dataset for three consecutive ep^16^ochs to avoid overfitting (**Supplementary Fig.1**). Evaluation of the model on the testing dataset demonstrated that SpikeHunter was accurate and sensitive, achieving an F1-score of 0.990, precision of 0.994, recall of 0.987, specificity of 1.000, a Mathew’s correlation coefficient (MCC) of 0.990 and the area under the precision-recall curve of 0.997 (**Fig.1c**). Based on these results, we concluded that SpikeHunter is an effective tool for tailspike protein identification and utilized it in subsequent analyses.

### Identification of 231,956 prophage-encoded tailspike proteins from the genomes of five common pathogens

A total of 787,566 genomes from *Escherichia coli*, *Pseudomonas aeruginosa, Klebsiella*, *Acinetobacter,* and *Salmonella* in the NCBI Pathogen Detection database were analyzed to predict prophages, prophage-encoded tailspike proteins, and bacterial serotypes. Prophages were predicted in 99.4% (783,032) of the genomes, resulting in a total of 8,423,494 prophage genomes across the entire dataset (**Fig.2a**). The SpikeHunter model was used to identify tailspike proteins within the prophages, resulting in the identification of 231,956 tailspike proteins in 228,055 prophages, representing 17,932 vOTUs (**Supplementary Table 1**). Despite the relatively small fraction of prophage genomes with a predicted tailspike protein (2.7%), 25.7% of the bacterial genomes analyzed contained at least one prophage genome with a predicted tailspike protein. Typically only one tailspike protein was observed in each prophage genome, but 1.67% prophages (3814 out of 228,055) were found to encode two or more tailspike proteins (**Fig.2a**). In the most extreme case, a single prophage from a *K. pneumoniae* (GCA_003037395.1) genome contained 14 predicted and was similar to the ϕKp24 jumbo phage, which is known for its highly-branched tail structure and expanded host range^17^ (**Supplementary Fig.2**). Overall, 208,900 prophages with tailspike proteins were identified in 182,937 bacterial genomes that also had a serotype prediction, providing good coverage across the species and facilitating the subsequent multi-species analysis of tailspike protein-serotype associations.

**Fig.2.**
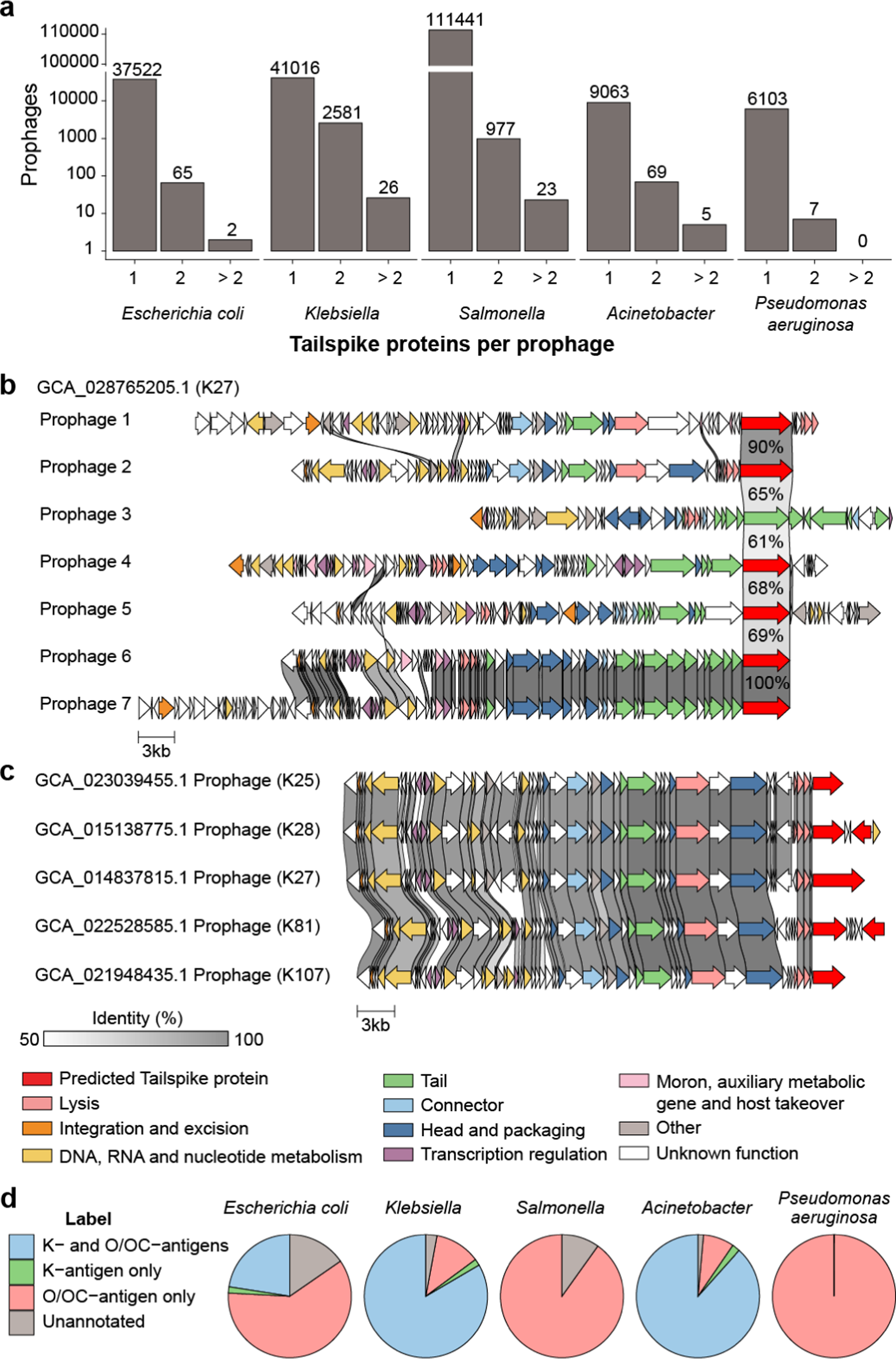
Identification of tailspike proteins and serotypes. a, Number of prophages detected in the genomes of each pathogen. b, Prophages detected in a *Klebsiella* genome (GCA_028765205.1) with the same tailspike protein clustered at 60% identity. c, Similar prophages with different tailspike proteins detected in multiple *Klebsiella* genomes. For panels b and c the amino acid similarity between genes is shown by the shaded region between the genes and the tailspike proteins are colored in red. d, Predicted serotypes for genomes of each of the pathogens. The relatively small number of genomes with K-antigen predictions for the *E. coli* can be partially attributed to the lack of Group 4 K-antigen prediction which would be functionally redundant with the strain O-antigen typing.

While most vOTUs (15,022 out of 16,535, 90.85%) only contained one type of tailspike protein (at 60 % identity), multiple examples (1,513 out of 16,535, 9.15%) of vOTUs composed of prophages with different tailspike proteins were found. In one example, a vOTU associated with *E. coli* was composed of prophages with 24 distinct tailspike proteins associated with distinct predicted serotypes, including a mix of K and O antigens (**Supplementary Fig.3**). Similarly, a *Klebsiella-*associated vOTU was identified where the five prophages each had a distinct tailspike protein and were associated with different K-antigen serotypes (**Fig.2c)**. The differences in tailspike protein and serotype associations for these otherwise similar prophages provides further evidence of the importance of the tailspike protein in driving host specificity. Similar tailspike proteins were found in distinct vOTUs from the same genomes (**Fig.2b**). Despite the difference in genomic content and organization between these vOTUs, their shared bacterial host further corroborates the link between the tailspike proteins and serotypes.

The distribution of serotypes in relation to tailspike proteins was then investigated. Bacterial serotypes were predicted for each of the bacterial genomes resulting in 182,937 bacterial genomes that had a predicted serotype and at least one prophage with an identified tailspike protein (**Fig.2d**). The tailspike proteins were subjected to hierarchical clustering analysis based on 30%, 60%, and 95% identity thresholds, and their relationship with the corresponding host serotypes of their (vOTUs) was examined (**Supplementary Table 2**). While the specific outcomes varied across different clusters, the predominant serotype associated with each cluster at the 30% identity level generally exhibited consistency with its descendant clusters at the 60% identity level. However, certain exceptions were observed. For instance, the cluster labeled cl30_19 in *Klebsiella*, which emerged at the 30% identity level, displayed high diversity within its descendant cluster at the 60% identity level (**Supplementary Fig.4**). Consequently, an identity threshold of 60% was proposed for all tailspike proteins to strike a reasonable balance between cluster purity and reducing redundancy in the results (**Supplementary Fig.5**).

### Prophage-encoded tailspike proteins are reliable indicators of bacterial serotypes

In the overall dataset, there were 1,180 unique tailspike protein to serotype associations identified, of which 715 (60.59%) were found to be strong associations (**Fig.3**). A majority of these strong associations were found from the *E. coli* (**Fig.3a**) and *Klebsiella* (**Fig.3b**) genomes, but multiple strongly associated tailspike-serotype pairs were still predicted for *Salmonella* (**Fig.3c**), *Acinetobacter* (**Supplementary Fig.6**), *and P. aeruginosa* (**Supplementary Fig.6**) providing coverage across all five taxa. These strongly associated tailspike-serotype pairs consisted of 681 unique tailspike clusters and 322 unique serotypes (127 K-antigens and 195 O/OC-antigens). Most of the strongly associated tailspike proteins were associated with only one serotype (99.56%) (**Fig.3**), suggesting that the association between tailspike proteins and serotype is consistent across different prophages.

**Fig.3.**
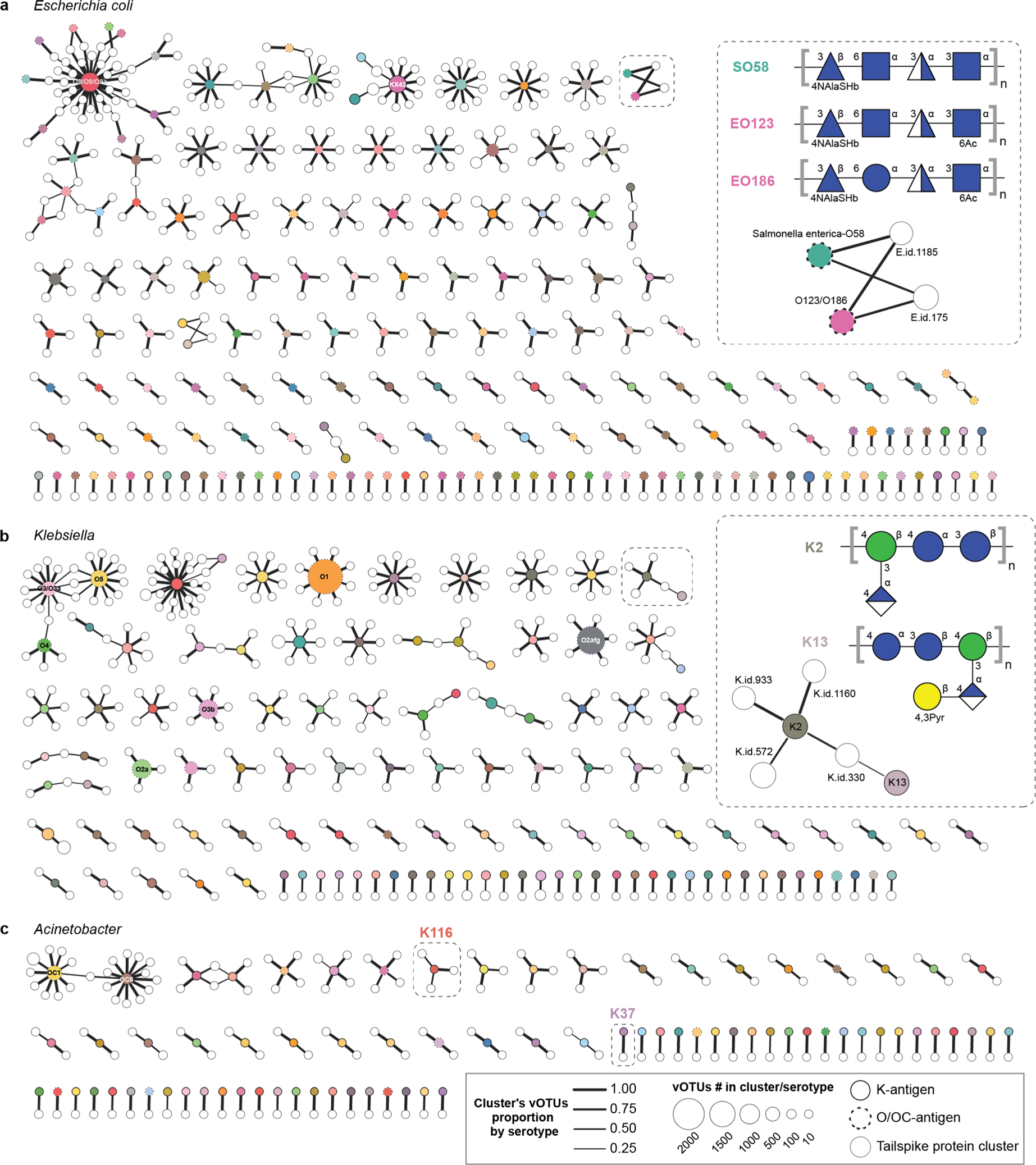
Association of tailspike protein clusters with bacterial serotypes. Networks showing the strongly associated serotypes (colored circles) with tailspike protein clusters (white circles). The size of the circle indicates the number of serotypes or tailspike proteins. Serotype circles with solid outlines represent K-antigens and circles with dashed outlines represent O/OC antigens. Width of the lines indicate the fraction of vOTUs containing that tailspike protein that support the association. Panels show the tailspike-serotypes associations for *E. coli* (a), *Klebsiella* (b), and *Acinetobacter* (c). Insets in panels A and B show the surface glycan structures associated with example mixed serotype clusters.

The association between prophage tailspike proteins and bacterial serotypes cannot be attributed to phylogenetic relatedness alone. The loci encoding bacterial surface glycans, which determine serotypes, are known to undergo frequent horizontal gene transfer^18^ polyphyletic distribution of serotypes. For instance, the serotype O81 in *E. coli* is distributed among multiple clades, and the tailspike protein cluster cl60_897 is found to strictly follow the clades that encompass the O81 serotype. Another example pertains to the K57 serotype in *Klebsiella*, where the associated tailspike cluster cl60_351 is present in three out of the four main K57 clades (**Supplementary Fig.7**). These findings suggest that the distribution of tailspike proteins aligns with serotypes rather than the phylogeny of the bacterial hosts.

Tailspike protein clusters that were strongly associated with multiple serotypes may be indicators of associations with other glycans or shared glycan structures between the serotypes. A few instances were observed in *E. coli* and *Klebsiella* where one tailspike protein was associated with two differently annotated serotypes. While glycan structural information for many of the serotypes is not available, examples were found for some of these components where the glycan antigen structures were similar. One tailspike protein was found to be associated with the *E. coli* O123 and O186 serotypes and the *S. enterica* O58 serotype, which have all been found to have a shared glycan backbone structure composed of 4-(N-acetylalanyl)amido-4,6-dideoxyglucose, N-acetyl-D-glucosamine, 2,5-dideoxy-2-(3-hydroxybutyramido)-glucose, and N-acetyl-D-glucosamine^19^ (**Fig.3a**). Similarly a tailspike protein was strongly associated with the K2 and K13 serotypes in *Klebsiella* which both share a backbone structure of two D-glucose units and a D-mannose unit^20^ (**Fig.3b**) and strains with these serotypes have been observed to be infected by the same phages^21^. These examples provide further evidence that tailspike proteins are associated with specific glycans structures and can provide valuable information about their associated bacterial host serotypes. Larger components containing links between multiple tailspike proteins and serotypes were observed for each of the five taxa (**Fig.3**). While some of these associations may be biologically relevant, these links are likely the result of either unpredicted serotypes that are common to the genomes and associated with the tailspike proteins or due to other factors like the association of the tailspike proteins with glycans that are not K or O/OC-antigens.

The associations between the tailspike proteins and serotypes also agreed with a well-documented case of successful phage therapy, corroborating the predictive power of this approach. A successful example of phage therapy being used can be seen in the example of the phage AbTP3phi1, which was effective against a multidrug-resistant *A. baumannii* infection with the K116 serotype^22, 23^. When compared to the set of tailspike proteins identified in this study, the tailspike protein from the AbTP3phi1 phage (UNI74976.1) was found to be homologous to two clusters of tailspike proteins associated with the Acinetobacter K37 (38.0% amino acid identity) and K116 (37.4% amino acid identity) serotypes (**Fig.3d**), two of the rarer serotypes only being predicted for 2 and 22 of the 7,001 Acinetobacter genomes analyzed respectively. The K116 serotype has been suggested to be a hybrid of genes from the Acinetobacter K37 and K14 serotype-specific loci, and the polysaccharide structures were found to have similar structures^24^. This result demonstrates the predictive capacity of tailspike protein-serotype associations, which stems from the comprehensive and large-scale analysis of the dataset. The knowledge pertaining to tailspike proteins can subsequently be employed to evaluate the viability of specifically targeting a phage towards a particular serotype, offering a resource to optimize the search process in phage therapy and rational phage engineering.

### Highly similar tailspike proteins are found across species with closely related surface glycans

Highly similar tailspike proteins (above 95% identity) were found in prophages from genomes of different species. Nearly all of the tailspike protein clusters (at 95% identity) were composed of tailspike proteins found in prophages from just one species (*i.e.*, only *E. coli*, only *S. enterica*, etc.). However a small fraction were made up of a mixture of tailspike proteins from prophages found in *E. coli* and *S. enterica* (26/41) or *E. coli* and *K. pneumoniae* (15/41) genomes (**Supplementary Table 3**). These mixed-species tailspike clusters were relatively rare but were all supported by at least two genomes from each species, reducing the chance that these were due to contamination or errors in the genomes.

The identification of similar tailspike proteins in *E. coli* and *Klebsiella* genomes with the same K-antigen suggests that the relationship between serotype and tailspike protein serves as a pivotal determinant of the host range. Instances of horizontal gene transfer of the K-antigen specific loci between *E. coli* and *Klebsiella* strains have been documented numerous times^25^ providing a possible route for phages to become associated with new host species. When examining the tailspike protein clusters that contained tailspike from prophages from different species, it was found that, despite their shared tailspike proteins, the prophages were distinct from each other (**Fig.4**). When examined in the context of their host serotypes it was found that these different phages from different species were associated with the same or similar serotypes. The E. coli genomes containing these tailspike proteins were predicted to have serotypes that were either the same as the *Klebsiella* strains in the cluster in the instance of the K47 (**Fig.4a**) and K63 (**Fig.4b**) serotypes or possessed a similar glycan backbone in the instance of the K13 and K2 serotypes (**Fig.4b and Fig.3b, Supplementary Table 3**).

**Fig.4.**
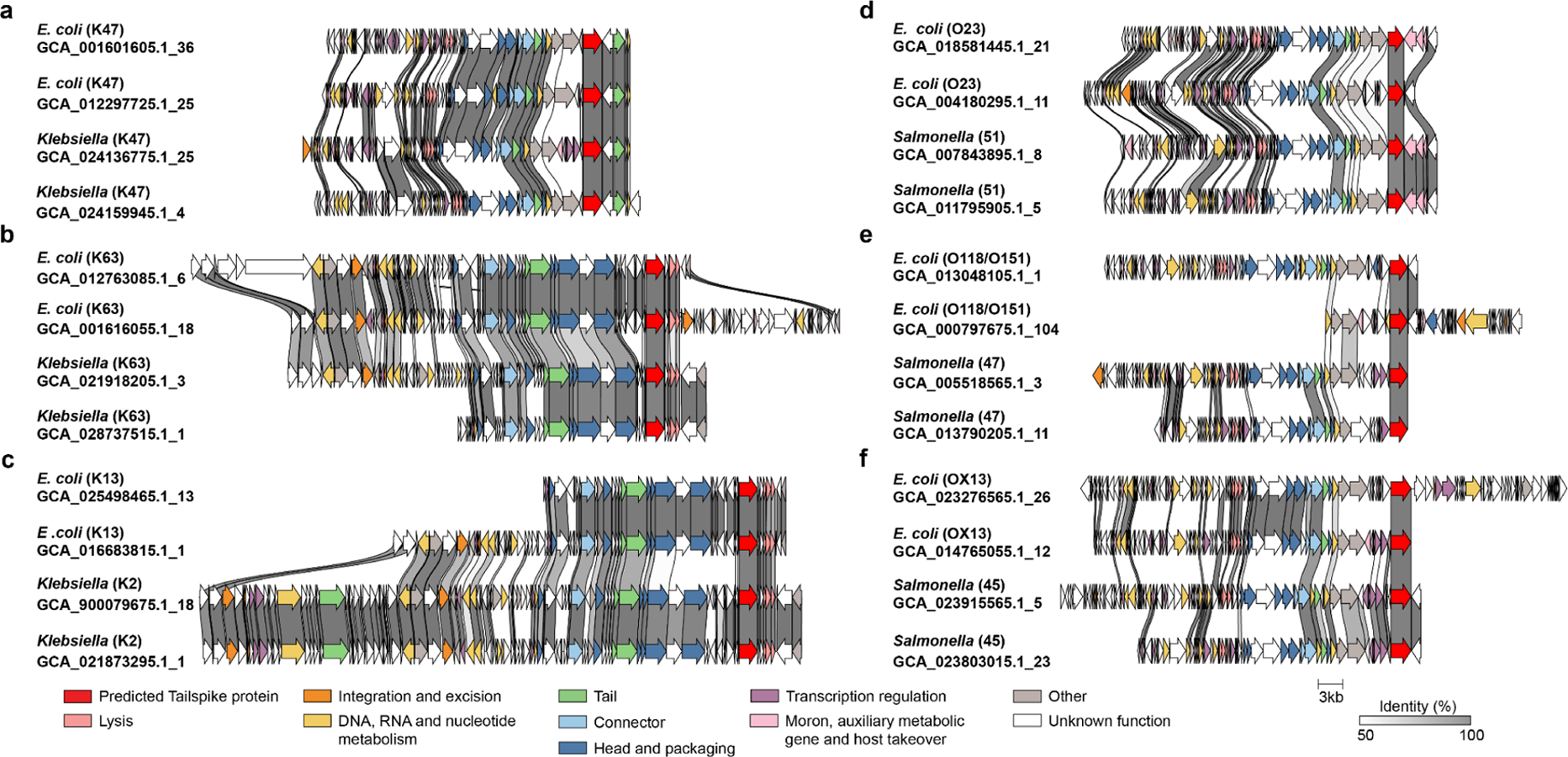
Shared TSPs among different phages. Prophage genomes from different species with the same tailspike protein are shown for *E. coli* and *K. pneumoniae* (a-c), and *E. coli* and *S. enterica* (e-f). Genes are colored based on their annotations using Pharokka with tailspike proteins being outlined in red and the amino acid identity of the genes is shown as the shaded regions between genes.

Examples of similar tailspike proteins were also observed in *E. coli* and *Salmonella* genomes suggesting that they shared similar O antigens. *E. coli* and *Salmonella* strains have also been found to possess similar O-antigen serotypes due to their evolutionary relatedness and horizontal gene transfer^26, 27^. One tailspike protein cluster was found in prophages associated with *E. coli* genomes predicted to have the O23 serotype and *Salmonella* genomes predicted to have the O51 serotype (**Fig.4d**), which have been found to have highly similar glycan backbone structures consisting with slightly different side groups (**Supplementary Figure 8a**)^28^. Additionally, the *E. coli* serotypes O118 and O151 are similar to the *Salmonella* serotype O47, differing by only the linkages between N-acetyl-beta-D-glucosamine and ribitol sugar residues^29^, suggesting that the tailspike proteins observed in the phages associated with these bacteria could interact with both glycan types (**Fig.4e, Supplementary Figure 8b**). Another set of phages were found to be associated with the *Salmonella* O45 serotype and *E. coli* OX13 serotype (**Fig.4f**). While the structure for the OX13 surface glycan antigen is not known, the O-antigen gene loci of the *E. coli* and *Salmonella* genomes were found to be highly similar in gene orders and gene identities with over 50% identity observed in 13 out of 14 gene pairs, suggesting a likely relationship between the glycan structures (**Supplementary Figure 8c**).

These multi-species tailspike protein to serotype associations provide further evidence of the strong associations of the tailspike proteins to surface glycan antigens. Cases of likely horizontal gene transfer present an interesting mechanism by which bacteria can become susceptible to types of phages and how the host range of phages could expand. Additionally, the association of tailspike proteins with distinct, but structurally similar glycan structures provides evidence that a limited degree of tailspike protein cross-reactivity with glycans may play a role in the expanded host range of some phages.

### Domain swapping in tailspike proteins

Extensive domain swapping between tailspike protein was observed in all five species. Tailspike proteins are commonly classified into an N-terminal head domain and a C-terminal domain (**Fig.5a**), with the C-terminal domain encompassing a beta-helix domain that is responsible for both receptor binding and depolymerase activity^13, 30^. Previous studies have reported the occurrence of naturally-occurring domain swapping in phages based on the presence of highly similar domains^13, 31^. Leveraging the large dataset of tailspike protein, a comprehensive assessment of domain swapping was performed. To convincingly identify these potential swaps, one of the domains is required to remain within a highly similar cluster (at 95% identity), while the other domain is placed in a distinct cluster, even if its identity threshold is significantly lower (at 30% identity). Evidence of N- and C-terminal domain swaps was observed in all five species, with a majority of the observed swaps being in tailspike proteins associated with *E. coli* or *Klebsiella* genomes. This observation can be attributed to the higher abundance of tailspike proteins identified in these two species, emphasizing the advantages offered by large datasets.

**Fig.5.**
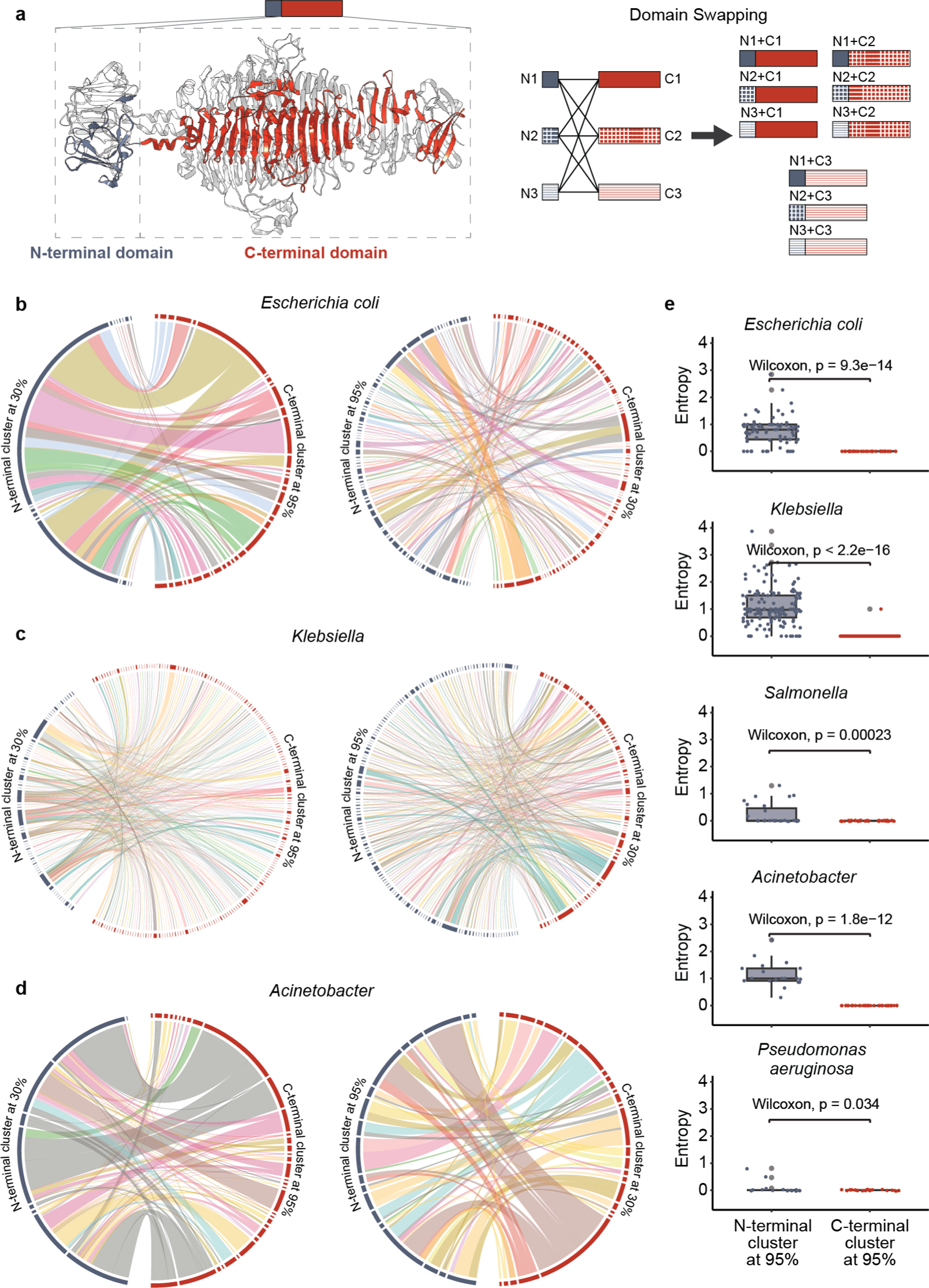
Domain swapping in tailspike proteins. a, Example domain partitioning of tailspike protein (PDB: 2XC1) and diagram illustrating domain swapping between hypothetical tailspike proteins. Putative domain-swapping in tailspike proteins from *E. coli* (b), *Klebsiella* (c), and *Acinetobacter* (d) are shown as connections between the N-terminal and C-terminal domains clustered at different amino acid identities. Bands connecting N and C-terminal domains are colored based on the serotype of the associated bacterial genomes. e, Boxplots comparing the entropy of serotypes observed in bacterial genomes with N and C-terminal domains clustered at 95% from proteins with potential domain-swapping. Significance of N and C-terminal entropy difference based on a Wilcoxon rank sum test is shown above bracket on each plot.

For the tailspike proteins that displayed putative domain swaps, an additional investigation was carried out to explore the relationship between their associated serotypes and the clustering of their N-terminal and C-terminal domains (**Fig.5b-d, Supplementary Figure 9**). Within the 95% identity cluster, it was observed that all tailspike proteins sharing the same C-terminal cluster were associated with the same serotype. However, this consistent serotype association was not observed across all N-terminal clusters. At the 30% identity level, the C-terminal domain clusters continued to exhibit a notably stronger association with serotypes compared to the N-terminal domain clusters. For example, at the 30% identity level, the C-terminal domain cluster, C_cl30_83, containing 274 vOTUs in *Klebsiella* exclusively corresponded to two serotype, whereas a N-terminal domain cluster, N_cl30_15, with a similar amount of vOTUs (215) in *Klebsiella* at the same identity level originated from a tailspike protein associated with twenty serotypes. Analysis of the serotype cluster entropy associated with each N-terminal and C-terminal domain corroborated these trends, revealing that C-terminal domains were characterized by significantly lower levels of serotype diversity compared to N-terminal domains in all of the species (**Fig.5e**).

## Discussion

Through this large-scale, multi-species analysis, we have demonstrated the strong association of tailspike proteins and bacterial serotypes, highlighted cases of inter-species serotype associations and domain swapping, and provided a rich resource that can guide the future application of phages in biomedicine. By applying protein language models, we have developed a sensitive approach for the detection of tailspike proteins, something that has been challenging due to the sequence diversity seen in these proteins. Our use of prophage-encoded tailspike proteins provides a way to examine confirmed tailspike-serotype associations across thousands of genomes without the need for isolating phages and bacterial strains. These factors have resulted in a rich dataset that captures phage and serotype diversity and can serve as a resource for future phage research.

Phage therapy has been proposed as an ideal tool for combating antibiotic resistance^8^ and suppressing disease-related members of the microbiome^32, 33^, but the application of phage therapy has consistently been limited by the ability to identify phages that would be effective against specific bacterial infections^8^. The similarity of the tailspike protein cluster associated with *Acinetobacter* K116 to the tailspike protein from phage AbTP3phi1 that was used in a successful phage therapy application^22^ demonstrates the utility of this data in guiding the selection of phages in the future. These results also demonstrate that domain swapping is common in tailspike proteins and the modular nature of tailspike proteins has allowed them to be engineered to alter their specificity^34^. Because isolating effective natural phages can be time-consuming, engineering targeted phages could be an effective strategy for combating new antimicrobial-resistant bacteria. This process and the application of tailspike proteins as antimicrobial compounds can be significantly enhanced by these results leading to more efficient and effective biomedical applications of phages.

Tailspike proteins also have a promising future as molecular tools in the glycosciences. The study of bacteriophages and phage-host interactions has led to the discovery of specialized polymerases^35^, ligases^35^, restriction enzymes^36^ and the CRISPR-Cas9^37^ system, which has revolutionized molecular biology. The glycan-targeting nature of tailspike proteins makes them ideal candidates as new tools to study glycans. Glycan profiling is inherently difficult due to the structural complexity, heterogeneity, lack of templates, and the limited availability of high-throughput analytical methods^38–40^. The depolymerase activity exhibited by tailspike proteins presents the potential for their utilization analogous to restriction enzymes in DNA digestion, facilitating the breakdown of polysaccharides into smaller fragments amenable to analysis, such as linkage analysis. Moreover, tailspike proteins have demonstrated utility in the development of pathogen biosensors reliant on the recognition of distinct glycans^41–43^, thus suggesting the possibility of expanding their functionality to establish novel glycan typing microarrays.

This approach to identifying tailspike protein associations has some disadvantages that highlight persistent gaps in the field. First, serotyping prediction tools are primarily focused on common human pathogens, limiting these results to these species of interest and underscoring the need for advanced serotype prediction tools that can be generalized to other species. Additionally, various other types of glycans that were not considered in this analysis are presented on the cell surface and have been shown to be receptors for some phages, including enterobacterial common antigen^2^ and cellulose^44^ non-tailspike phage receptor binding proteins also play roles in binding to bacterial cell surface glycans and proteins^45^ are important considerations for future studies investigating phage-host interactions. These limitations highlight gaps in the understanding of both bacterial serotypes and phage biology that will be important aspects of future research. Expanding this work to include free phages and to account for other bacterial species will enhance the associations and provide important context to the results and facilitate their application in biomedical contexts. Overall this study provides an essential foundation for the future study of bacteriophage host range and the future use of phages and tailspike proteins in a variety of fields.

## Methods

### Training and validation data

A total of 3659 bacteriophage genomes were downloaded from the INPHARED database^16^ (v1.7). This collection of proteins was split into tailspike and non-tailspike phage protein datasets. A possible tailspike protein dataset was generated based on keyword searches and comparisons to other annotated viral proteins in the NCBI nr database using BlastP, PDB using SCOP^46^, and the viral ortholog databases, PHROG^47^, pVOG^48^, ViPhOG^49^, eggNOG viral ortholog groups^50^, and VOGDB (https://vogdb.org/), using HMMER. Proteins that were annotated as tailspike, tail fiber, or receptor binding were included in the candidate tailspike dataset along with proteins whose top hit was annotated as tailspike proteins in other databases. The set of candidate tailspike proteins were then clustered at 70% identity using CD-HIT^51, 52^ and their structures were predicted using AlphaFold^53^. The structures were then manually curated to identify a final set of 1,913 tailspike proteins based on the presence of the distinctive beta-helix receptor-binding domain. The remaining 201,994 non-tailspike proteins (excluding those classified as part of the ‘unknown’ category in INPHARED) were included in the non-tailspike dataset.

### Model architecture

The SpikeHunter model was developed using the PyTorch framework^54^. First, the phage sequences are first tokenized and transformed into numerical vectors using the *batch_converter* function in the ESM python package^55^. The sequences are then embedded as 1280 length representations using a pre-trained transformer protein language model ESM-2 (esm2_t33_650M_UR50D)^15^. The sequence representations are fed into a three-fully-connected layer network with 1280, 568 and 2 nodes, respectively. The output from the last layer is converted into a probability representing each sequence being a tailspike protein or not with a softmax activation function. The SpikeHunter model and code is available on GitHub (https://github.com/nlm-irp-jianglab/SpikeHunter).

### Training and validation of the deep learning model

To train and validate the SpikeHunter model the manually curated set of tailspike proteins was first clustered into 19,399 clusters at 30% identity using CD-HIT^52^. The sequences were then divided into training, validation, and testing datasets in a ratio of 3:1:1 using the *StratifiedGroupKFold* function in the *Scikit-learn* python package^56^, resulting in a training set of 125,085 proteins (comprising 1,106 positive samples and 123,979 negative samples belonging to 11,638 clusters), a validation set of 38,584 proteins (comprising 361 positive samples and 38,223 negative samples belonging to 3,878 clusters) and a testing set of 40,238 proteins (comprising 446 positive samples and 39,792 negative samples belonging to 3,883 clusters). The model training was performed with the cross-entropy loss function and the Pytorch implementation of the Adam optimizer, with the parameters of the ESM-2 model frozen. The training was halted when the model’s performance on the validation dataset did not improve for three consecutive epochs. The model with the lowest validation loss was then used for testing and prediction.

### Identification of tailspike proteins in bacterial genomes

A total of 787,566 genomes of five common pathogens, *E. coli*, *K. pneumoniae*, *P. aeruginosa*, *S. enterica,* and *A. baumannii*, were downloaded from the NCBI Pathogen Detection database (https://www.ncbi.nlm.nih.gov/pathogens, downloaded on 2nd April, 2023. Prophage regions were predicted in the pathogen genomes using VIBRANT (version 1.2.0) with default parameters to identify phages within the bacterial genomes. The protein sequences of the phages were then extracted and SpikeHunter was used to classify them as either tailspike or non-tailspike proteins. Only proteins with greater than 50% probability of being a tailspike protein were considered to be positive hits for tailspike proteins.

### Serotyping of the bacteria genomes

Serotype prediction for the five analyzed microbial species was performed using a variety of species specific tools. O-antigen serotypes were predicted for *E. coli* using ECTyper^57^, SerotypeFinder^58^ and fastKaptive^18^. Serotypes were grouped into serogroups according to Iguchi *et al.,*^59^. K-antigen prediction for *E. coli* genomes was done using fastKaptive^18^. *K. pneumoniae* K- and O-antigen serotypes were inferred using Kaptive^58^ and fastKaptive^18^. *A. baumannii* K-antigen, and OC serotypes were predicted using Kaptive^58^. *P. aeruginosa* O-antigen serotypes were predicted using PAst^60^. O-antigen serotypes were predicted for *S. enterica* using SeqSero2^61^ and fastKaptive^18^. All tools were run using default settings. Results from Kaptive with match confidence scores of “None” and results from fastKaptive with best match coverages lower than 90 were excluded from the downstream analysis. To facilitate the comparison of serotypes between species, shared serotypes were merged and consistently named for the set of *K. pneumoniae* and *E. coli* K-antigens and the set of *E. coli* and *S. enterica* O-antigens. Common serotypes between *K. pneumoniae* and *E. coli* were identified by predicting serotypes in the *K. pneumoniae* genome using the fastKaptive tool. The predicted *K. pneumoniae* serotypes from Kaptive and fastKaptive were then used to map the shared serotypes between the two species. The same procedure was followed for identifying common serotypes between *S. enterica* and *E. coli*, but with the *S. enterica* genomes being annotated using SeqSero2 and fastKaptive to provide the mapping to the *E. coli* genomes.

### Associating tailspike proteins with serotypes

Tailspike protein sequences were hierarchically clustered at 30%, 40%, 50%, 60%, 70%, 80%, 85%, 90% and 95% identities using cd-hit v.4.8.1^62^. The tailspike protein clusters at different levels were associated with the vOTUs they were found in and the serotypes of the genomes that the vOTUs were in. When encountering multiple instances of the same 95% tailspike protein cluster, vOTU, and serotype within the dataset, only one instance was retained for subsequent analysis to eliminate redundancy. An overall network of tailspike protein clusters at 60% identity and serotypes was then generated with the links between them being determined by the vOTUs. Associations between the tailspike protein clusters at 60% identity and serotypes were classified into three categories, “highly confident”, “confident”, and “uncertain”. “Highly confident” associations were tailspike to serotype pairs that were supported by at least 90% of the vOTUs containing that 60% tailspike protein cluster for tailspike proteins with more than four vOTU connections or that were supported by all vOTUs for tailspike protein clusters with less than or equal to 4 vOTUs. “Confident” associations were tailspike to serotype pairs that were supported by 10% to 90% of the vOTUs with the tailspike protein cluster and the remaining associations were classified as “uncertain”. More information regarding the assignment of serotypes to a tailspike protein cluster can be found within the code hosted on GitHub (https://github.com/nlm-irp-jianglab/TSP_paper).

### Identification of domain swapping between tailspike proteins

An all-by-all BlastP^63^ search was performed to compare all non-redundant tailspike protein sequences, and the hits were filtered using an e-value cutoff of 1e-8, identity threshold of 60%, and a coverage range of 10% to 90% of the sequence. Candidate domain-swapping was found by identifying proteins that aligned to at least two other proteins that cumulatively covered at least 90% of the original query sequence. Breakpoints for the N and C-terminal domains were determined by computing the average alignment start and end positions from all of the alignments to each query tailspike protein. The domain sequences were then extracted and clustered using psi-cd-hit v.4.8.1 at 30% and 95% identity thresholds^62^. Domain-swapping was only considered plausible if they were validated by multiple proteins with the same domain at 95% identity while having different versions of the other domain at 30%. Circlize v0.4.15 was used to visualize the N/C-terminal domain-swapping for each species^64^.

## Author contributions

Conceptualization and Analysis: YY, KD, WY, and XJ. Methodology: YY, TC, LX, XJ. Writing: YY, KD, XJ.

## Acknowledgments

This work utilized the computational resources of the NIH HPC Biowulf cluster. (http://hpc.nih.gov). We thank Dr. Audrey Burnim at NLM/NIH for help with visualizing protein structures and Hui Yi from CenHTRO at the University of Georgia for her assistance with the statistical approaches used in the study.

## Data and materials availability

The authors confirm that the data supporting the findings of this study are available within the article and its supplementary materials.

