## Supplementary Figures for "Deciphering Phage-Host Specificity Based on the Association of Phage Depolymerases and Bacterial Surface Glycan with Deep Learning"

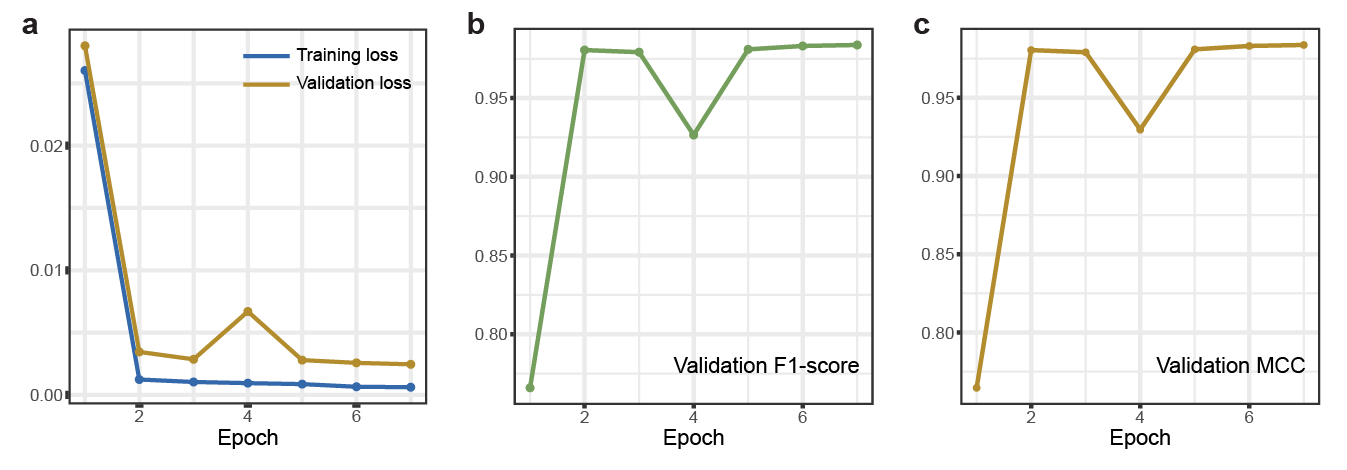

**Supplementary Figure 1. SpikeHunter training and validation loss.** The training loss and validation loss (a), validation F1-scores (b), and validation MCC (c) in each epoch during the training of SpikeHunter.

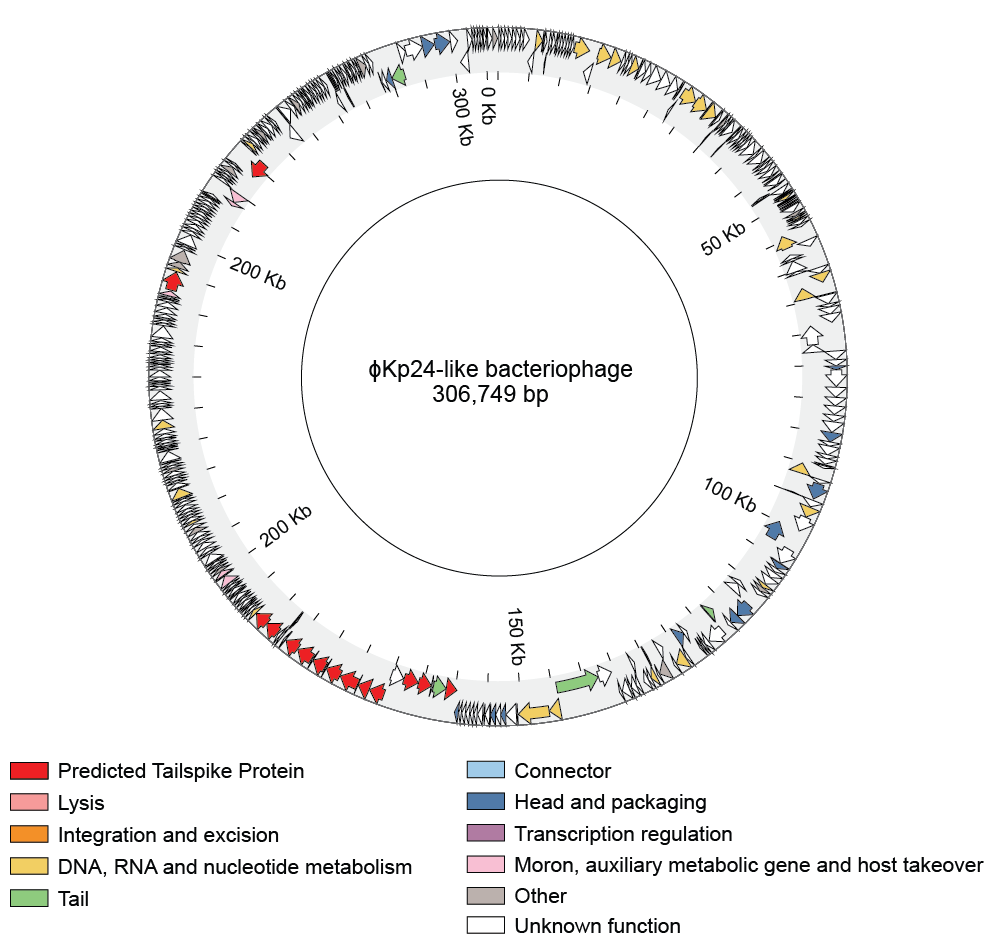

**Supplementary Figure 2.** **ϕKp24-like jumbo prophage.** Circular diagram of a ϕKp24-like prophage genome identified in a *K. pneumoniae* genome (GCA_003037395.1). Predicted genes are colored based on Pharokka annotations and predicted tailspike proteins are colored red.

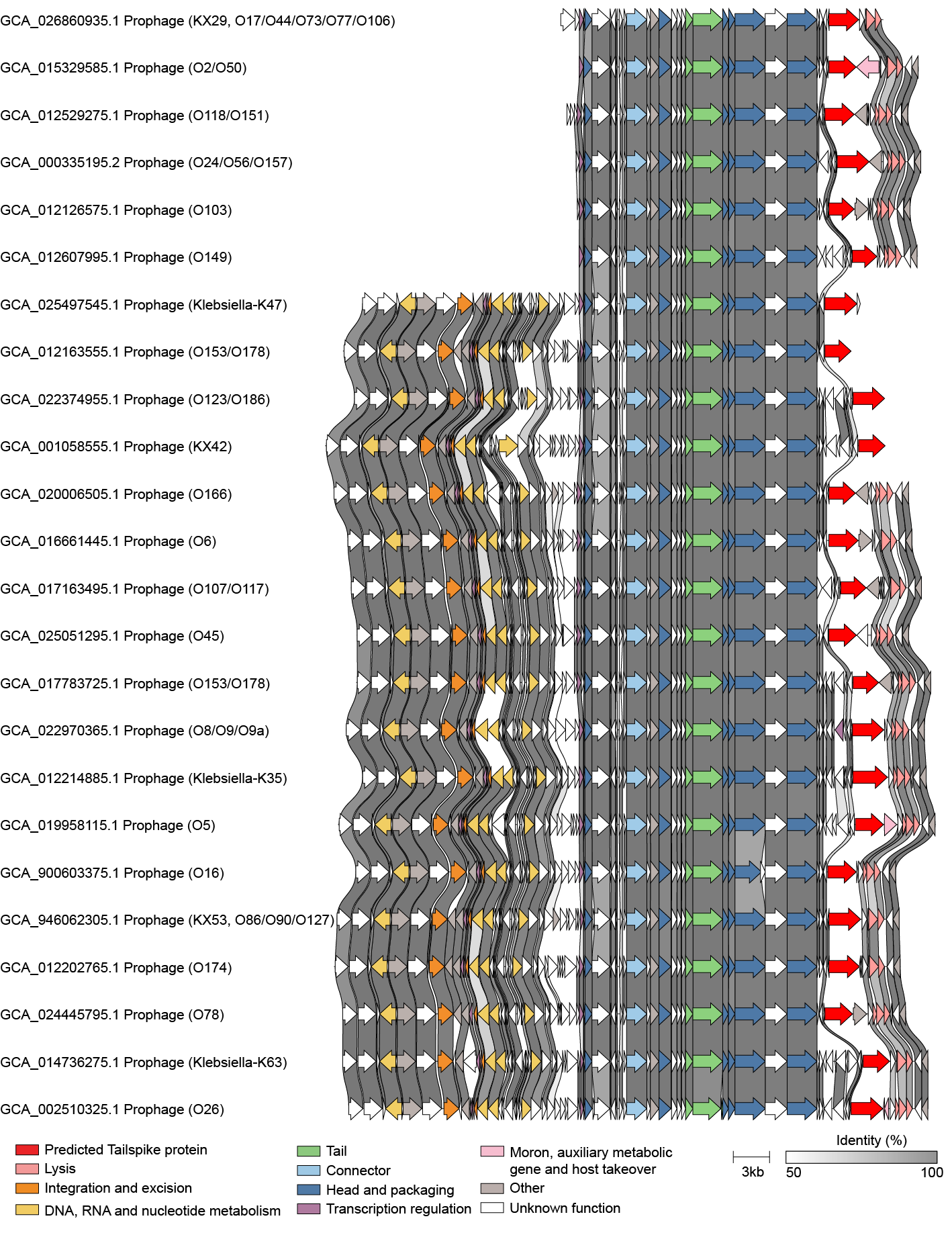

**Supplementary Figure 3.** ***E. coli* phages with 24 serotypes. Diagram of 24 similar *E. coli* from the same vOTU that have distinct tailspike proteins and are associated with distinct serotypes.** Genes are colored based on Pharokka annotations and tailspike proteins are colored red. Shaded regions between genomes indicate the amino acid identity between the genes. The genomes and their predicted serotypes are labeled on the left of each phage genome.

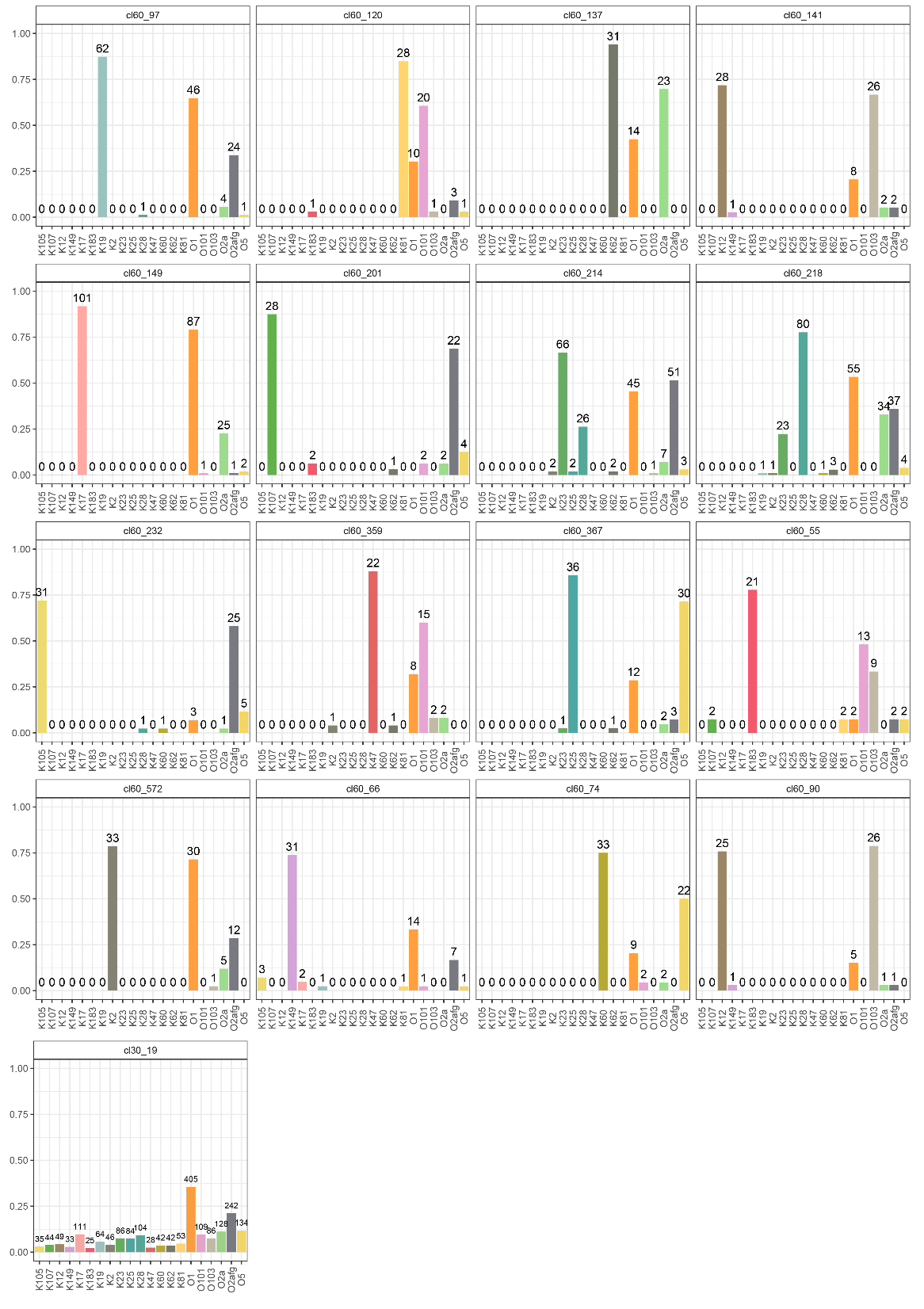

**Supplementary Figure 4.** **High serotype diversity within a tailspike protein cluster.** Plots showing the fraction of genomes associated within the cluster that serotype. Some genomes can be associated with both K and O antigens meaning that total fractions for a cluster can add up to more than 1. The 30% identity cluster is shown on the left, with its composite 60% identity clusters being shown in the plots in the right four columns. The number of tailspike proteins within the cluster that are associated with each serotype are listed above each bar.

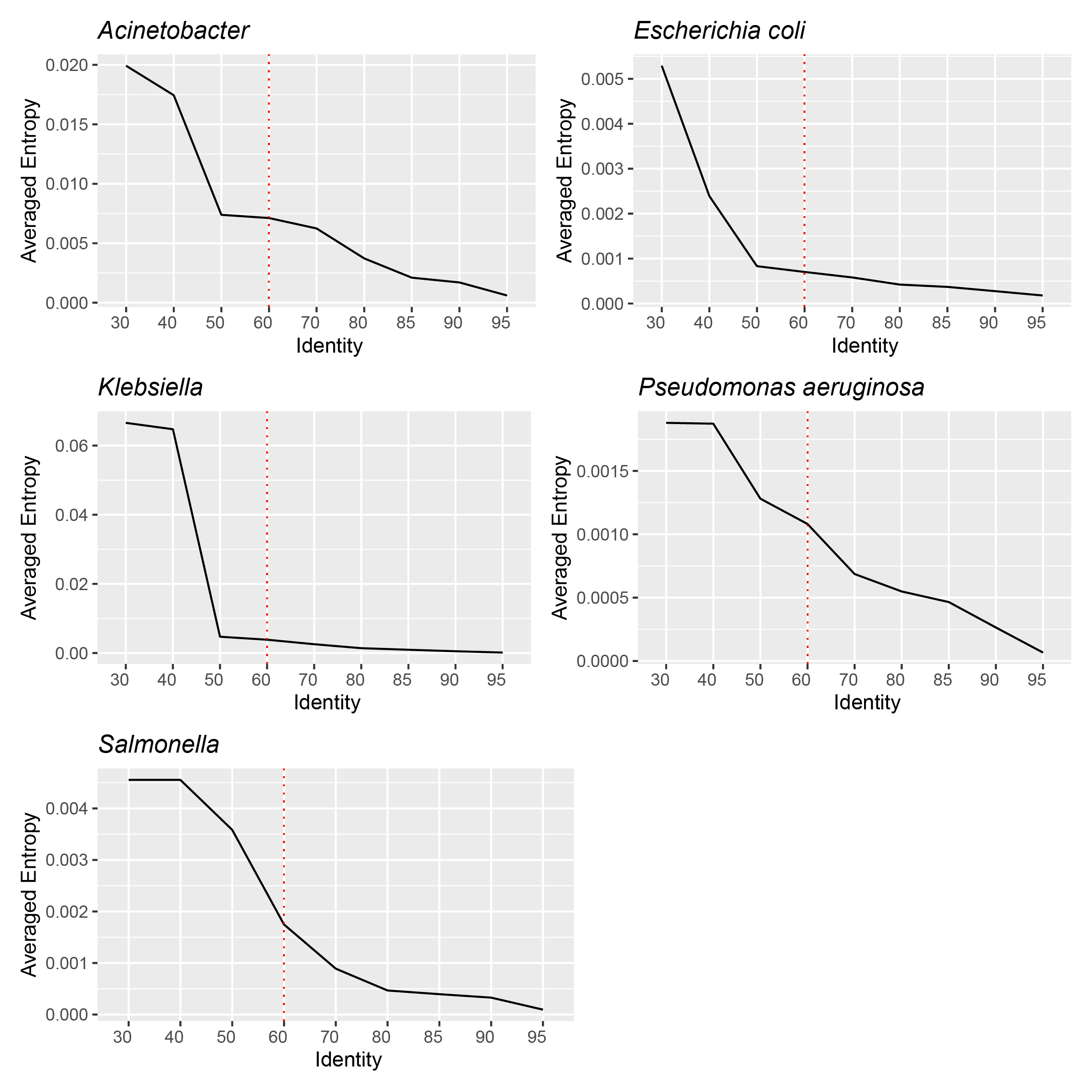

**Supplementary Figure 5.** **Purity of serotypes associated with 60% identity tailspike protein clusters.** The x-axis of the plots shows the percent identity used to cluster the tailspike proteins and the y-axis shows the corresponding average entropy of the serotypes associated with the clusters at that identity level. The red dotted line marks the 60% clustering threshold used for downstream tailspike protein to serotype analyses.

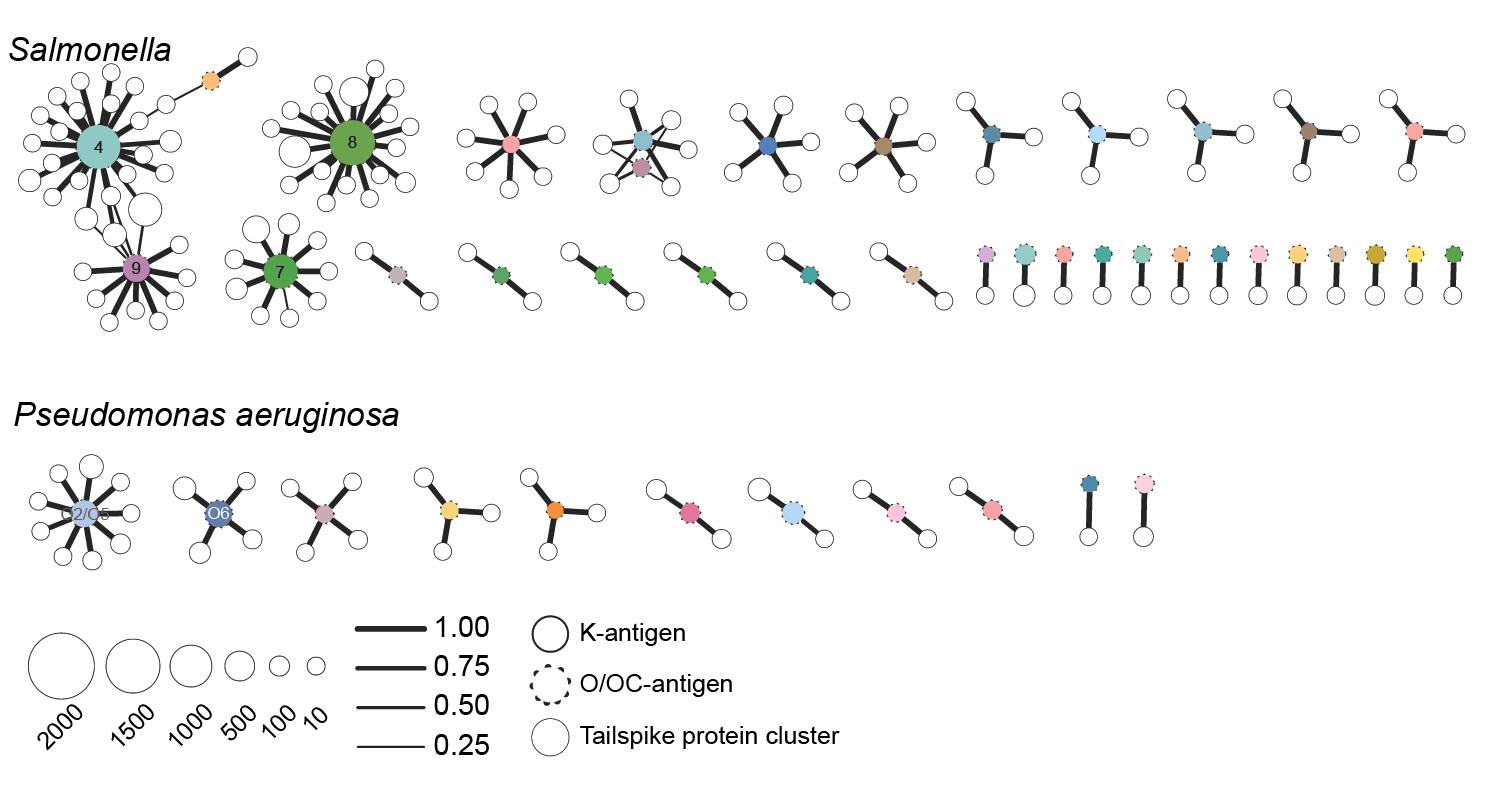

**Supplementary Figure 6.** **Tailspike protein to serotype networks for *Acinetobacter* and *P. aeruginosa*.** Networks showing the strongly associated serotypes (colored circles) with tailspike protein clusters (white circles). The size of the circle indicates the number of serotypes or tailspike proteins. Serotype circles with solid outlines represent K-antigens and circles with dashed outlines represent O/OC antigens. Width of the lines indicate the fraction of vOTUs containing that tailspike protein that support the association.

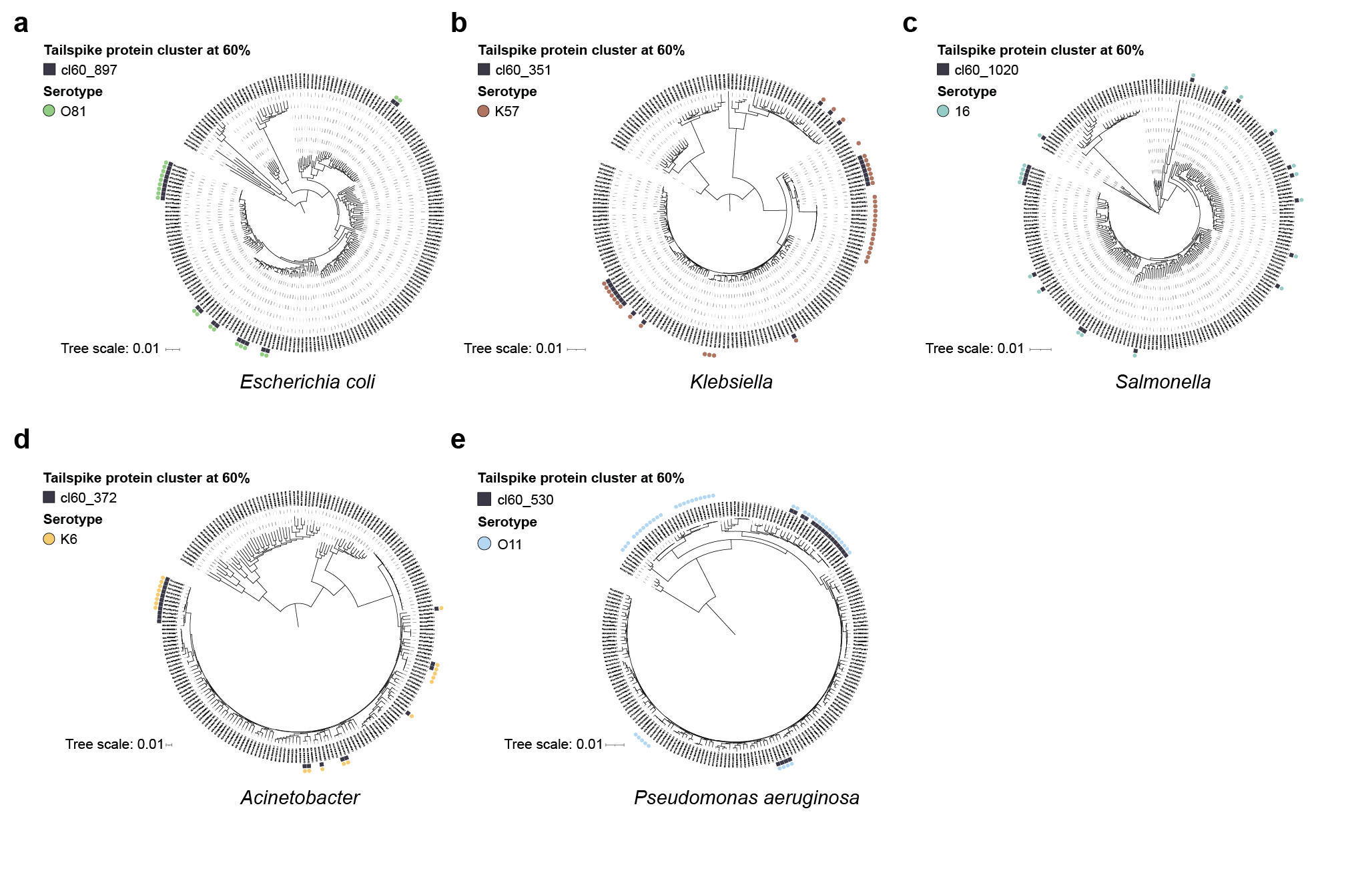

**Supplementary Figure 7.** **Example phylogenetic distributions of tailspike proteins and serotypes.** The phylogenetic distribution of a tailspike protein and serotype are shown for *E. coli* (a), *Klebsiella* (b), *Salmonella* (c), *Acinetobacter* (d), and *P. aeruginosa* (e).

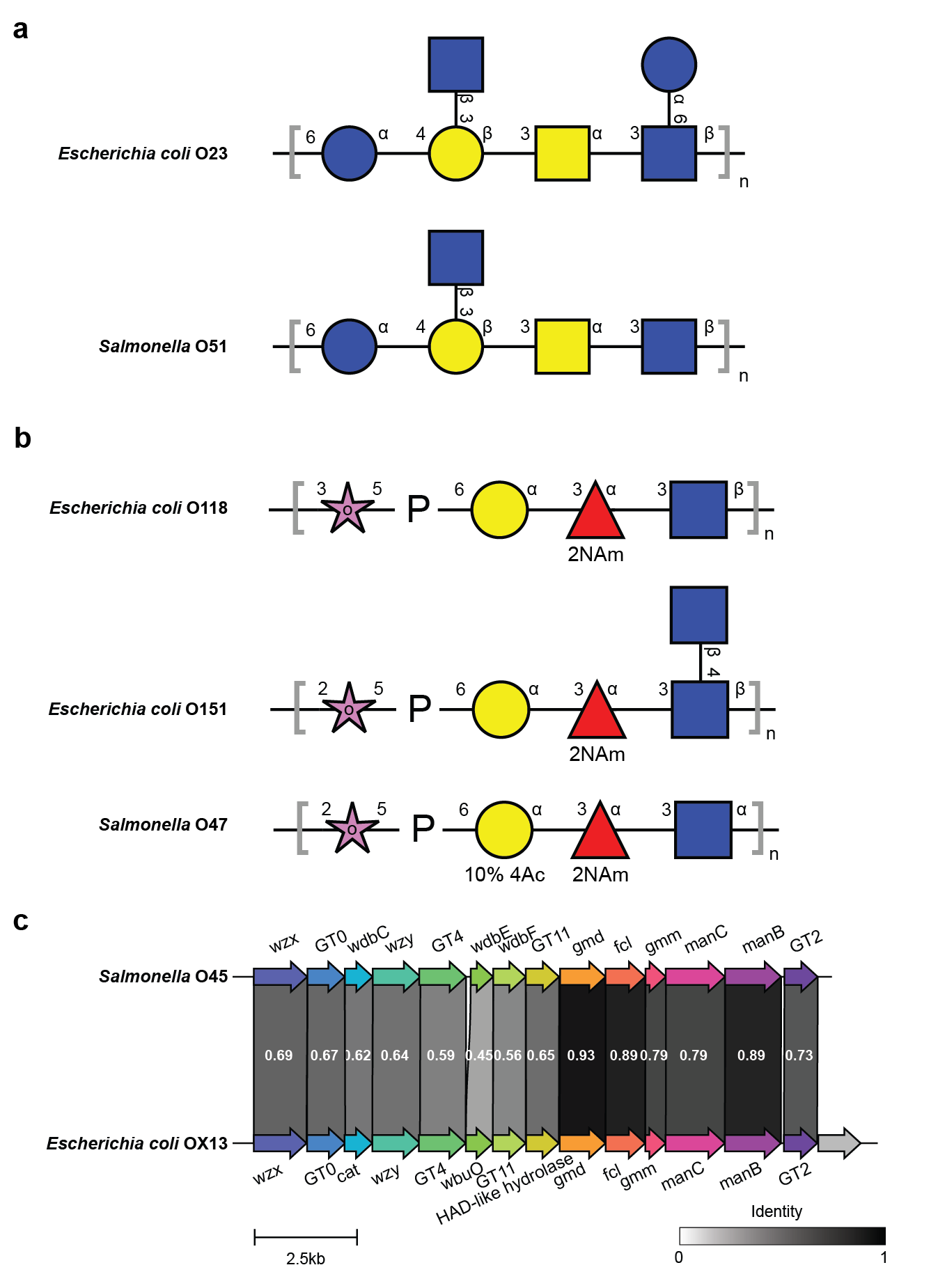

**Supplementary Figure 8.** **Support for tailspike proteins associated with *E. coli* and *Salmonella*.** Glycan antigen backbone structure similarity is shown in panels a and b. Panel c shows the similarity in gene clusters between the OX13 genetic loci (KP710591.1) and *Salmonella* O45 genetic loci (JX975332.1). The shaded region between the genes indicates the amino acid identity of the genes.

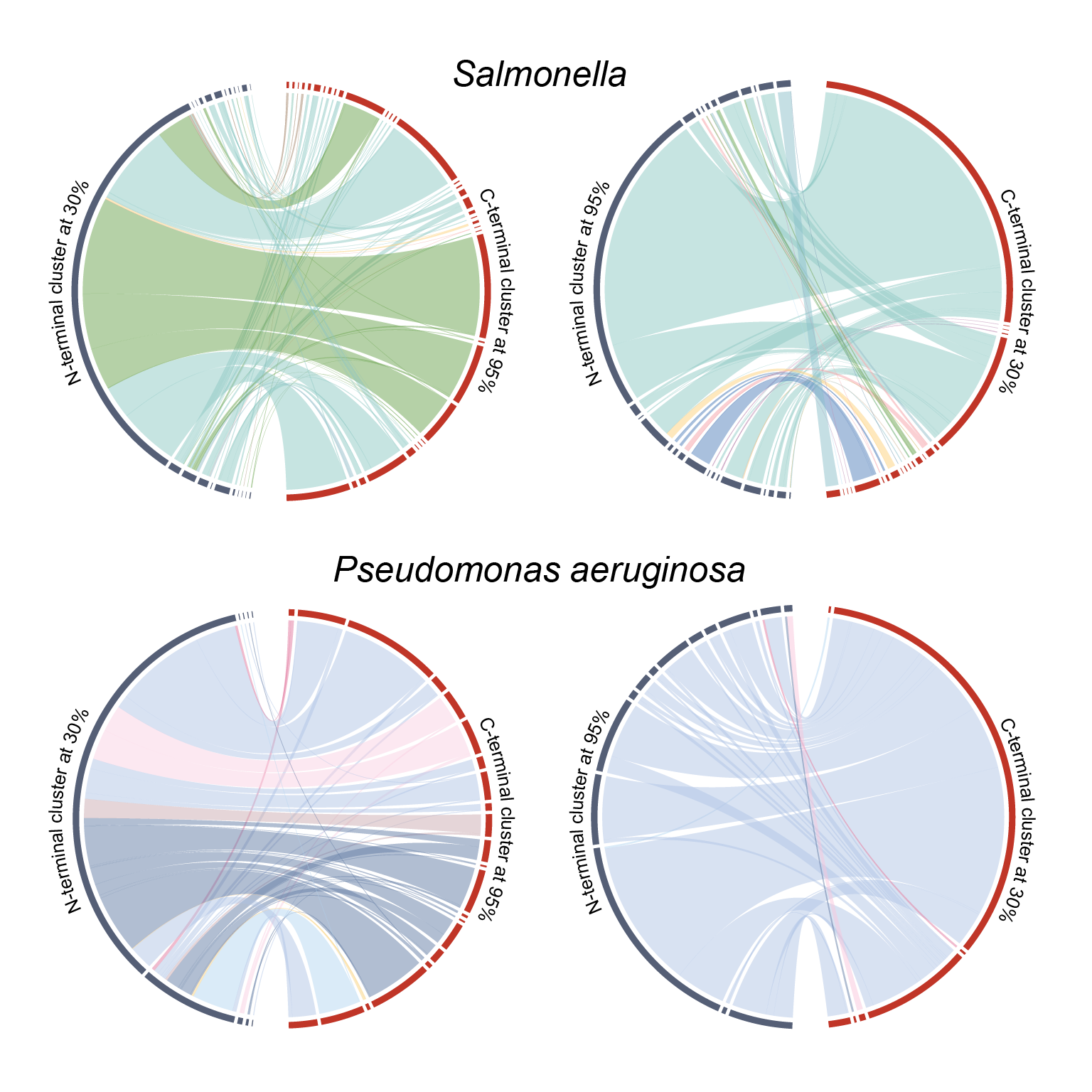

**Supplementary Figure 9.** ***Acinetobacter* and *P. aeruginosa* domain swapping.** Putative domain swapping is shown as connections between the N-terminal and C-terminal domains clustered at different amino acid identities. Bands connecting N and C-terminal domains are colored based on the serotype of the associated bacterial genomes.
