## Supplementary Information for "Deciphering Phage-Host Specificity Based on the Association of Phage Depolymerases and Bacterial Surface Glycan with Deep Learning"

**Supplementary Tables**

**Supplementary Table 1.** **Tailspike protein clusters at different identities.**

**Supplementary Table 2. Tailspike protein cluster to serotype associations.**

**Supplementary Table 3.** **Cross-species tailspike protein associations.**

**Supplementary Figures**

**Supplementary Figure 1. SpikeHunter training and validation loss.**

**Supplementary Figure 2.** **ϕKp24-like jumbo prophage.**

**Supplementary Figure 3.** ***E. coli* phages with 24 serotypes. Diagram of 24 similar *E. coli* from the same vOTU that have distinct tailspike proteins and are associated with distinct serotypes.**

**Supplementary Figure 4.** **High serotype diversity within a tailspike protein cluster.**

**Supplementary Figure 5.** **Purity of serotypes associated with 60% identity tailspike protein clusters.**

**Supplementary Figure 6.** **Tailspike protein to serotype networks for *Acinetobacter* and *P. aeruginosa*.**

**Supplementary Figure 7.** **Example phylogenetic distributions of tailspike proteins and serotypes.**

**Supplementary Figure 8.** **Support for tailspike proteins associated with *E. coli* and *Salmonella*.**

**Supplementary Figure 9.** ***Acinetobacter* and *P. aeruginosa* domain swapping.**
